# Senescent cells and the dynamics of aging

**DOI:** 10.1101/470500

**Authors:** Omer Karin, Amit Agrawal, Ziv Porat, Valery Krizhanovsky, Uri Alon

**Author notes:** equal contribution (Omer Karin, Amit Agrawal).

## Abstract

A causal factor in mammalian aging is the accumulation of senescent cells (SnCs) with age. SnCs cause chronic inflammation, and removing SnCs decelerates aging in mice. Despite their importance, however, the production and removal rates of SnCs are not known, and their connection to aging dynamics is unclear. Here we use longitudinal SnC measurements and SnC induction experiments to show that SnCs turn over rapidly in young mice, with a half-life of days, but slow their own removal rate to a half-life of weeks in old mice. This leads to a critical slowing-down that generates persistent SnC fluctuations. We further demonstrate that a mathematical model, in which death occurs when fluctuating SnC populations cross a threshold, quantitatively recapitulates the Gompertz law of survival curves in mice and humans. The concept of a causal factor for aging with rapid turnover which slows its own removal can go beyond SnCs to explain the effects of interventions that modulate lifespan in *Drosophila* and *C. elegans*, including survival-curve scaling and rapid effects of dietary shifts on mortality.

## Main Text

Senescent cells (SnCs) increasingly accumulate with age in mice and humans in many tissues^1–7^, due in part to DNA damage, damaged telomeres, and oxidative stress^5,8^. These cells, characterized by high levels of p16 and SA-β-Gal^5^, enter permanent cell cycle arrest, and secrete a characteristic profile of molecules including pro-inflammatory signals^9^ and factors that slow regeneration^9^ (Figure 1A). They have physiological roles in development, cancer prevention and wound healing^9–11^. However, as organisms age, accumulating levels of SnC cause chronic inflammation and increase the risk of many age-related disease including osteoarthritis, neurodegeneration, and atherosclerosis^12–24^.

**Figure 1.**
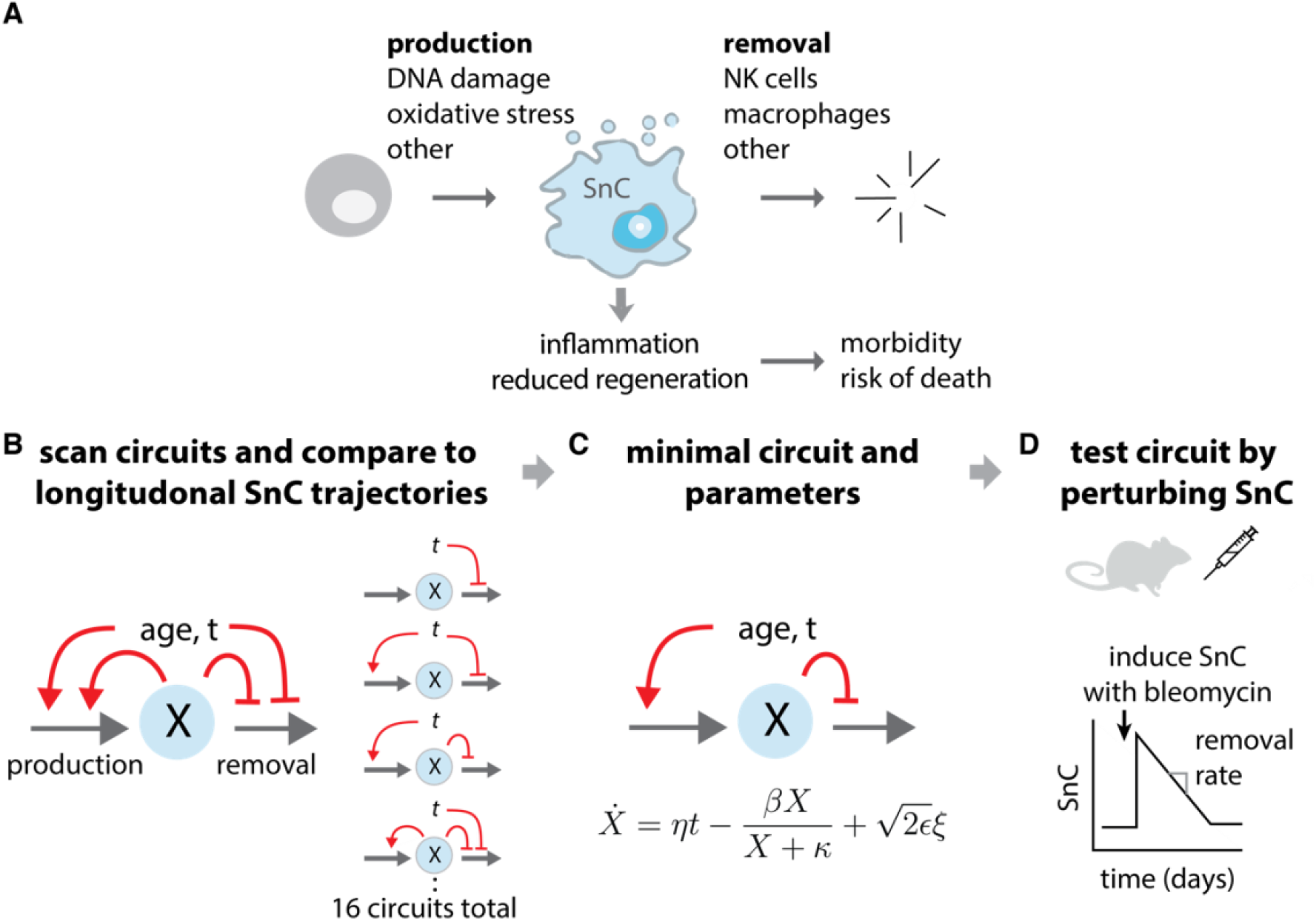
Approach for inferring SnC dynamics throughout adulthood. (A) Many processes, including DNA damage and developmental and paracrine signals, lead to SnC production. SnCs are cleared by immune mechanisms, and secrete factors that lead to morbidity and mortality. (BCD) We scanned a wide class of models for SnC dynamics, and compared them to longitudinal SnC data and direct SnC perturbation experiments to establish a minimal model for SnC stochastic dynamics and its rate constants. In the minimal model, η is the increase in SnC production rate with age, β is the removal rate, κ is the half-way saturation point for removal and ϵ is the noise amplitude.

Accumulation of SnCs is in fact known to be causal for aging in mice: continuous targeted elimination of whole-body SnCs increases mean lifespan by 25%, attenuates age-related deterioration of heart, kidney, and fat, delays cancer development^25^ and causes improvement in the above mentioned diseases.

These studies indicate that SnC abundance is an important causal variable in the aging process. Despite their importance, however, the production and removal rates of SnCs are unknown^9,26^. For example, it is unclear whether SnCs passively accumulate or if they are turned over rapidly, and if so, whether their half-life changes with age. Since turnover affects the ability of a system to respond to fluctuations, information about these rates is crucial in order to mathematically test ideas about the possible role of SnCs in the age-dependent variations in morbidity and mortality between individuals.

Here, we address this experimentally and theoretically. To understand the dynamics of SnCs, we scanned a wide class of mathematical models of SnC dynamics, and, as described below, compared these models to longitudinal SnC trajectories^1^ and direct SnC induction experiments in mice (Figure 1BCD). The models all describe SnC production and removal. They differ from one another in the way that production and removal rates are affected by age and by SnC abundance. The models describe all combinations of four possible mechanisms for accumulation of SnCs: (i) SnC production rate increases with age due to accumulation of mutations^27^, telomere damage and other factors that trigger cellular senescence^28^; (ii) SnCs catalyze their own production by paracrine and bystander effects^29^, (iii) SnC removal decreases with age due to age-related decline in immune surveillance functions^30^, and (iv) high SnC levels reduce their own removal rate. This reduction can be due to SnCs saturating immune surveillance mechanisms or due to SnC-related signaling or disruption of cellular architecture that interferes with removal. These four effects lead to 16 different circuits (Figure 1B) with all combinations of whether or not each of effects (i-iv) occur. Additionally, each of the 16 models includes parameters for basal production and removal. The models have rate constants that are currently uncharacterized.

To find which of the model mechanisms best describes SnC dynamics, and with which rate constants, we compared the models to longitudinal data on SnC abundance in mice collected by Burd et al^1^. SnC abundance was measured using a luciferase reporter for the expression of p16^INK4a^, a biomarker for senescent cells. Total body luminescence (TBL) was monitored every 8 weeks for 33 mice, from early age (8 weeks) to middle-late adulthood (80 weeks) (Figure 2A). We tested how well each model describes the longitudinal SnC trajectories by finding the maximum likelihood parameters for each of the 16 models, adjusting for number of parameters (Supplementary Section 1 and Section 2).

**Figure 2.**
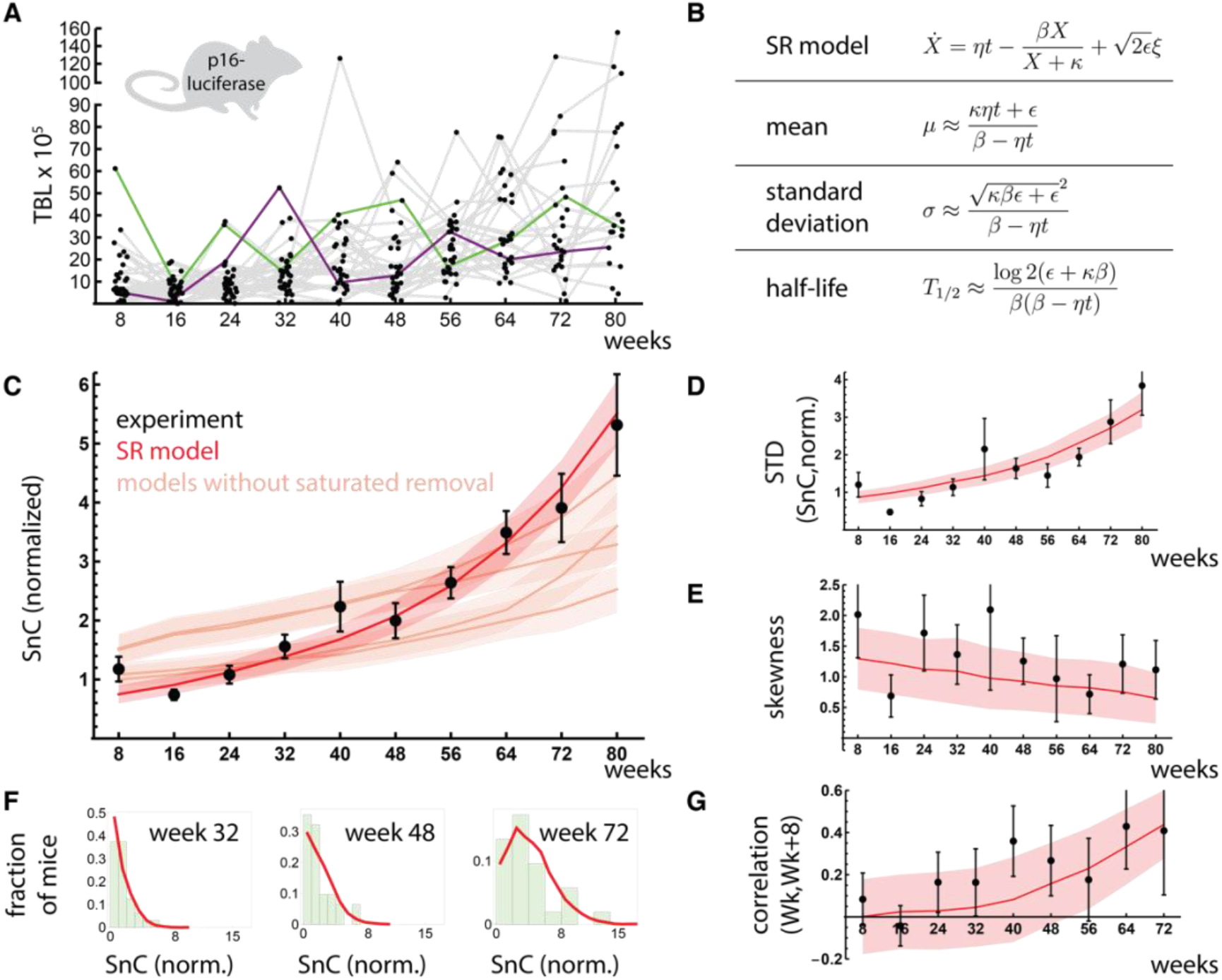
Saturated-removal (SR) model captures longitudinal SnC trajectories in mice. (A) Total body luminescence (TBL) of p16-luciferase in mice. Grey lines connect data from the same individual mice (green and purple lines are examples of individual trajectories). (B) SR model equations and their approximate analytical solutions. The SR model (red line) captures (C) the mean SnC abundance, (D) standard deviation of SnC abundance, (E) skewness and (F) shape of the distributions among equal-aged individuals, and (G) correlation between subsequent measurements on the same individuals. TBL was normalized to give a mean abundance of 1at young ages. Maximum likelihood parameters for the SR model are: *η* = 0.15 *day*^−1^*year*^−1^, *β* =0.27 *day*^−1^, *κ* = 1.1, *ϵ* = 0.14 *day*^−1^. Orange lines in (C): best-fit of models without saturation (Supplementary Section 1). Mean and standard error (shaded red, orange regions) are from bootstrapping.

A principle emerges from this comparison: in order to capture the longitudinal dynamics, the mechanism must have rapid turnover of SnCs on the timescale of a few days in young mice, and it also must include mechanism (iv), which we henceforth call ‘saturation of removal’. The simplest model that describes the data has only two interactions (Figure 2B): SnC production rate increases linearly with age, and SnCs slow down their own removal rate, similar to saturation of an enzyme by its substrate. We call this the saturating removal model (SR model), whose equation is given in Figure 2B.

The SR model captures the accelerating rise of mean SnC abundance with age in the longitudinal data (Figure 2C): as SnCs accumulate, they slow their own removal, leading to even higher SnC levels. The SR model also explains the increasing SnC variability between individuals which accelerates with age (Figure 2D), and the SnC distributions among equal-aged individuals (Figure 2E), which are skewed to the right (Figure 2F).

Importantly, the SR model captures the fact that SnC fluctuations become more persistent with age, as evidenced by an increasing correlation between subsequent measurements (Figure 2G, p<0.01): individuals with higher (or lower) than average SnC levels stay higher (or lower) for longer periods with age. This increased persistence is due to the effect of SnCs on their own removal rate. Models without mechanism iv (saturation of removal) show a poor overall fit (light-red lines in Figure 2C).

The maximum likelihood parameters of the SR model (listed in the caption of Figure 2) provide quantitative predictions for SnC half-lives: SnC turnover is rapid in young mice, with a half-life of about 5±1 days at 3 months of age; Turnover slows with age, so that SnC half-life is about 25±6 days at 22 months.

We tested these predictions using experiments in mice by inducing senescent cells and analyzing their dynamics. To induce senescence in mice lungs we used intra-tracheal bleomycin administration (Figure 3A), a DNA damaging agent that induces cellular senescence in the lung epithelium a few days after treatment^5,31^.

**Figure 3.**
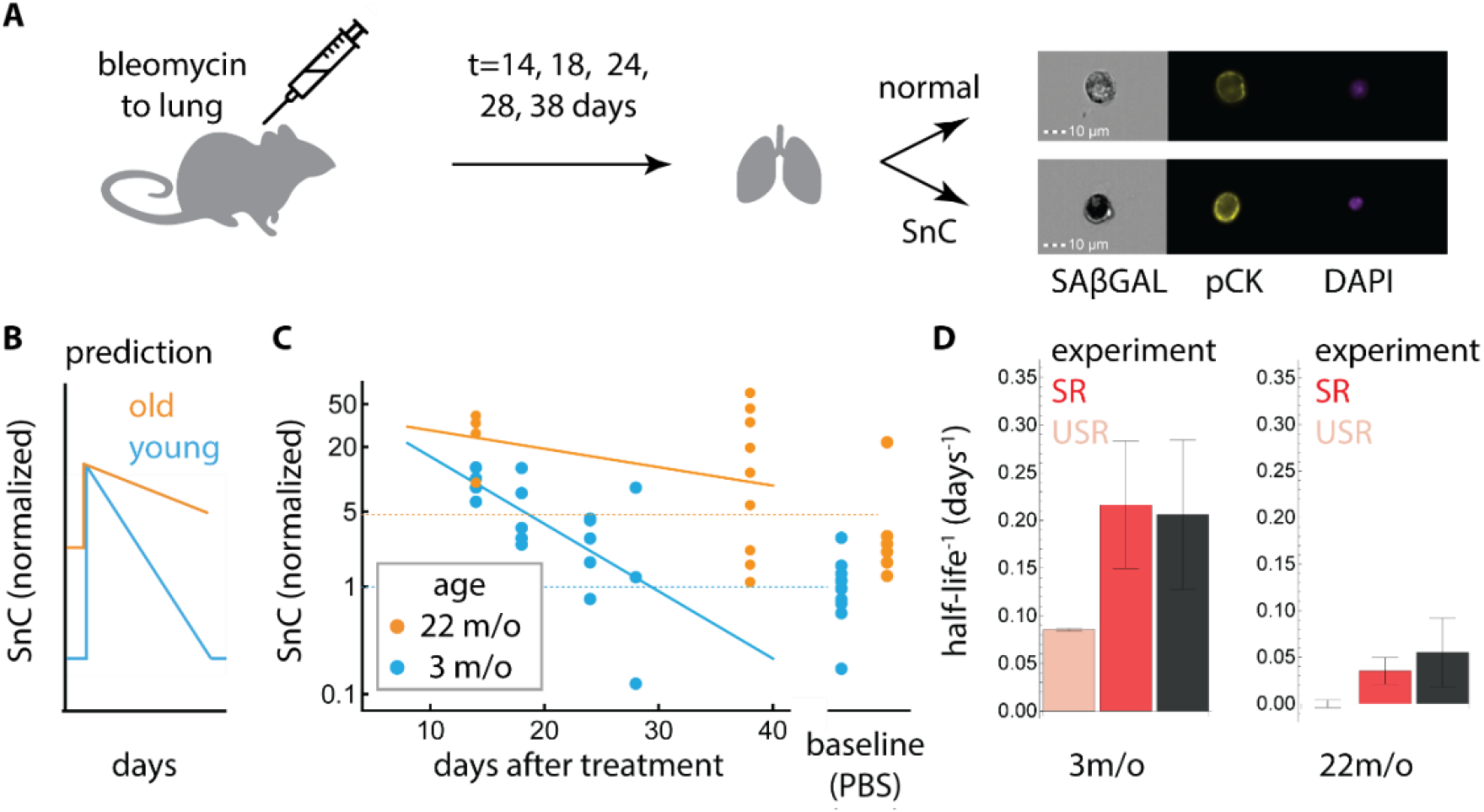
SnC half-life measurements in mice support SR model predictions. (A) Bleomycin or PBS was introduced by intratracheal installation to mice on day 0. Lungs were analyzed on the indicated days thereafter. Representative images of lung cells analyzed by imaging flow cytometry show how senescent epithelial cells were identified, using SA-β-Gal, Pan-Cytokeratin (pCK), and DAPI staining. SnC removal rate was estimated by log-linear fit. (B) The SR model predicts that SnCs rapidly return to baseline in young mice and that removal is slower in old mice. (C) Fraction of SnCs in mouse lungs after treatment with bleomycin (1.5U/Kg). In young mice, SnC levels return to baseline with a half-life of about 5 days. In old mice, baseline SnC levels are about 5-fold higher, and SnC removal rate is slower than in young mice (p=0.038). (D) SnC removal rates (halflife^-1^) for young and old mice (mean and SE, black) agree with the SR model predictions (red, mean and SE were calculated by bootstrapping, see Methods). The best-fit model without saturated removal (USR) shows a poor prediction (orange).

We quantified the fraction of senescent lung epithelial cells at different time points following bleomycin administration (Figure 3A) using imaging flow cytometry. Epithelial SnCs were defined as cells positive for a senescent cell marker (SA-β-Gal) and an epithelial marker (pan-Cytokeratin, pCK). This cell population was also HMGB1 nuclear negative, as expected in senescent cells^5,32^, and previously shown^5^ to be correspond to non-proliferative cells (negative Ki67 assay) (see Supplementary Section 3).

In 3-month-old mice, SnC levels decayed with a half-life of *τ* = 4.7 days (*τ*^−1^ = 0.21 ± 0.07 *days*^−1^) and reached their baseline level within less than a month (Figure 3BC), as predicted. SnC levels in young mice are thus in a rapid dynamic balance of production and removal.

To test the prediction that removal slows with age (Figure 3B), we performed the bleomycin treatment in old mice (22 month-old). In these mice, the baseline level of SnCs was about 5-fold higher than in young mice (Figure 3D). SnCs decayed with a half-life of *τ* = 18 days (*τ*^−1^ = 0.055 ± 0.035 *days*^−1^), slower than that of young mice as predicted (p=0.038, Figure 3B).

These turnover measurements quantitatively agreed with the predictions of the SR model (Figure 3D, Supplementary Section 4) with no additional fit. This agreement occurred despite the use of distinct SnC markers in the two data sets (SA-β-Gal in the bleomycin experiment vs. p16^INK4A^-luciferase in the longitudinal experiment), suggesting consistency between the measurement methods.

Our results suggest a core mechanism in which SnC production rate rises linearly with age, and SnCs slow their own removal (Supplementary Section 5). This slowdown of removal accelerates SnC accumulation with age. Slowdown of removal also amplifies fluctuations in SnC levels at old ages. This amplification, known as *critical slowing down*^33,34^, results in long-lasting differences among individuals at old ages. In other words, young mice have large spare removal capacity of SnC; old mice have much smaller spare removal capacity. This smaller removal capacity means that any addition of SnCs takes longer to remove, causing larger and more persistent variation in SnC levels among individuals (Figure 2G).

In the remainder of the paper, we use mathematical analysis to explore the implications of rapid SnC turnover and removal slowdown to address the question of variability in mortality. Mortality times vary even in inbred organisms raised in the same conditions, demonstrating a non-genetic component to mortality^35,36^. In many species, including mice and humans, risk of death rises exponentially with age, a relation known as the Gompertz law^37–39^, and decelerates at very old ages. The Gompertz law has no known explanation at the cellular level.

To connect SnC dynamics and mortality, we need to know the relationship between SnC abundance and risk of death^1^. The precise relationship is currently unknown. Clearly, SnC abundance is not the only cause for morbidity and mortality. It seems to be an important causal factor because removing SnCs from mice increases mean lifespan^25^, and adding SnCs to mice increases risk of death and causes age-related phenotypes^23^. We therefore explored the simple possibility that death can be modeled to occur when SnC abundance exceeds a threshold level *X_C_*, representing a collapse of an organ system or a tipping point such as sepsis (Figure 4A). Thus, death is modelled as a first-passage time process, when SnC cross X_C_. We use this assumption to illustrate our approach, because it provides analytically solvable results. We also show that other dependencies between risk of death and SnC abundance, such as sigmoidal functions with various degrees of steepness, provide similar conclusions.

**Figure 4.**
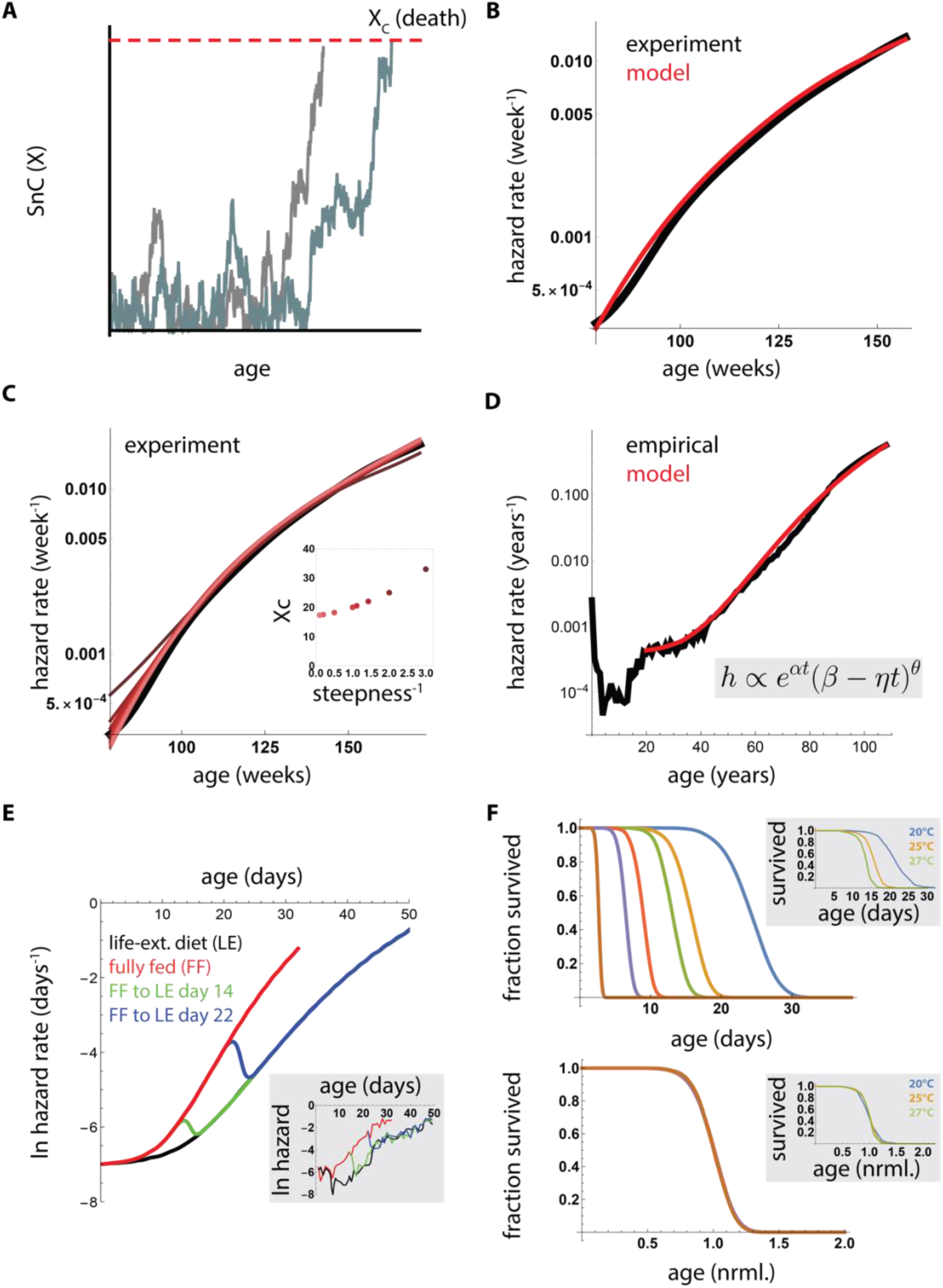
SR model can explain the variability in mortality between individuals. (A) To model the relation between risk of death and SnC levels, we assumed a simple threshold model where death occurs when SnC abundance exceeds a critical threshold X_c_ (B) Mouse mortality (C57BL/6J mice obtained from the Mouse Phenome Database^54^, black line) is well fit by the SR model (red line) with parameters consistent with the data of Figures 1 and 2, with death defined when SnC exceed a threshold (*η* = 0.084 *day*^−1^*year*^−1^, *β* = 0.15 *day*^−1^, *κ* = 0.5, *ϵ* = 0.16 *day*^−1^, *X_c_* = 17) (C) Similar results are obtained by assuming a more general sigmoidal dependency between SnC abundance X and risk of death: *h* = (1 + *e*^−*α*(*X–X_c_*)^)^−1^. Parameters are as in B, except that X_c_ is adjusted according to the steepness parameter α (*inset*). (D) The SR model with added age-independent extrinsic mortality of 0.4 · 10^−3^*year*^−1^ (red) matches human mortality statistics ^55^ (black). Inset: approximate analytical solution for the first passage time in the SR model shows the Gompertz law and deceleration at old ages. (E) SR model described rapid shifts in mortality when fully fed *Drosophila* transition to a lifespan-extending dietary intervention (LE), experimental data in inset ^45^ (*β* = 1 *hr*^−1^, *κ* = 1, and *ϵ* = 1 *hr*^−1^ *η* = 0.03*hr*^−1^*day*^−1^, *X_c_* = 15. LE modeled by a decrease in *η*, *η* = 0.02*hr*^−1^ *day*^−1^ (changes in other parameters lead to similar conclusions, see Supplementary Section 7). (F) Lifespan of *C. elegans* raised at different temperatures varies by an order of magnitude, but survival curves collapse on a single curve when time is scaled by mean lifespan (inset: data from ^35^). The SR model provides scaling for perturbations that affect *η*, but not other parameters (*β* = 1 *hr*^−1^, *κ* = 1, and *ϵ* = 1 *hr*^−1^, *η* = 0.07 *hr*^−1^ *day*^−1^, Supplementary Section 8).

The SR model analytically reproduces the Gompertz law, including the observed deceleration of mortality rates at old ages (Figure 4B-D, Supplementary Section 2). Notably, most models without both rapid turnover and slowdown of removal do not provide the Gompertz law (Supplementary Section 2). The SR model gives a good fit to the observed mouse mortality curve (Figure 4B-C, Supplementary Section 1) using parameters that agree with the present experimental half-life measurements and longitudinal SnC data (Supplementary Section 1). Thus, turnover of days in the young and weeks in the old provides SnC variation such that individuals cross the death threshold at different times, providing the observed mortality curves.

The SR model can describe the observed increase in mean lifespan of mice in experiments in which a fraction of SnCs are continually removed (Supplementary Section 6). More generally, the SR model can address the use of drugs that eliminate SnCs, known as senolytics^40^. To reduce toxicity concerns, it is important to establish regimes of low dose and large inter-dose spacing^41^. The model provides a rational basis for scheduling senolytic drug administrations. Specifically, treatment should start at old age, and can be as infrequent as the SnC turnover time (~month in old mice) and still be effective (Supplementary Section 6).

We also adapted our results from the mouse data to study human mortality curves. In humans, mortality has a large non-heritable component^42,43^. A good description of human mortality data, corrected for extrinsic mortality, is provided by the same parameters as in mice, except for a 60-fold slower increase in SnC production rate with age in the human parameter set (Figure 4D, Supplementary Section 7). This slower increase in SnC production rate can be due to improved DNA maintenance in humans compared to mice^44^. We conclude that the critical slowing-down described by the SR model provides a possible cellular mechanism for the variation in mortality between individuals.

The generality of the SR model suggests that it might also apply organisms where ageing may be driven by factors other than senescent cells, such as *Drosophila melanogaster* and *C. elegans*, in which lifespan variation is well-studied^35,45^. In these organisms, the present approach can be extended by considering X as a causal factor for aging, that accumulates with age and has SR-type dynamics^46^, namely turnover that is much more rapid than the lifetime, and self-slowing removal. One clue for the identity of such factors may be gene-expression variations in young organisms that correlate with individual lifespan^47–49^, and the actions of genes that modulate lifespan^39,50–53^.

Work in *C. elegans* and *Drosophila* provides constraints to test the SR model. For example, *Drosophila* shows rapid switches between hazard curves when transitioning between normal and lifespan-extending diets (Figure 4E, inset). These switches are well described by the SR model, due to its rapid turnover property (Figure 4E and Supplementary Section 8). The rapid turnover property entails that the level of *X* can adjust after a change in any of the parameters of the model. A model without rapid turnover could not explain these results.

We further tested whether the SR model can explain the scaling of survival curves for *C. elegans* under different life-extending genetic, environmental and diet perturbations^35^. These perturbations change lifespan by an order of magnitude, but the survival curves collapse on the same curve when age is scaled by mean lifespan (Figure 4F insets). We find that the SR model provides this scaling for perturbations that affect the accumulation rate *η* (Figure 4F). Interestingly, we predict loss of scaling when a perturbation affects other parameters such as removal rate *β* or noise *ϵ* (Supplementary Section 9), a prediction that may apply to exceptional perturbations in which scaling is not found such as the *eat-2* and *nuo-6* mutations (Supplementary Section 6). In all cases, scaling cannot be explained without rapid turnover. We conclude that the SR model of rapid turnover with critical-slowing down is a candidate explanation for scaling of survival curves in *C. elegans*.

In summary, we propose a framework for the dynamics of SnCs based on rapid turnover that slows with age. Bleomycin-induced SnC half-life is days in young mice and weeks in old mice, causing critical slowing down which greatly amplifies the differences between individual SnC levels at old age. We theoretically explore the implications of this slowdown in a model in which SnCs cause death when they exceed a threshold. The widening variation in SnC levels with age causes a mortality distribution that follows the Gompertz law of exponentially increasing risk of death. The mortality distribution of mice and humans is well-described by the SR model with the SnC half-lives measured here. Future work may test this proposed connection between SnC dynamics and mortality by experimentally measuring risk of death as a function of SnC abundance.

Our results suggest that treatments that remove senescent cells can have a double benefit: an immediate benefit from a reduced SnC load, and a longer-term benefit from increased SnC removal. Similarly, interventions that increase removal capacity, for example by augmenting the immune surveillance of SnC, are predicted to be an effective approach to reduce SnC levels and increase health-span.

## Acknowledgments

U.A. is the incumbent of the Abisch-Frenkel chair. O.K. is an Azrieli Fellow. This work was supported by grants to V.K. from the European Research Council under the European Union’s FP7 and H2020 Programs and the Israel Science Foundation.

