## appendix for "Senescent cells and the dynamics of aging"

### Methods

#### Stochastic model simulation.

Simulations of the stochastic models were performed by using the ItoProcess function of Mathematica (V11.0), with a step size of 1 day. Negative X values were avoided using a reflecting boundary condition at X=0. In simulations that included mortality, time of death was the first time-point where X exceeded $X_C$.

#### Model comparison to longitudinal p16INK-luciferase measurements.

We sought for each model the parameters that maximize the log-likelihood of the measured trajectories (Figure 1A). We calculated the log-likelihood of a model $m$ with parameters $\theta$ as follows. Let $X_{i,j}$ be the measured SnC level (SnC=TBL/9.63 to give SnC=1 for young mice) of mouse $j$ at time point $i$ (with $X_{0,j} = 0$). We denote by $Prob_{m,\theta,i}(a|b)$ the probability of reaching SnC level $a$ at time point $i$ given SnC level $b$ at time point i-1. We call such a step from *i-1* to *i* a sub-trajectory. We estimated this probability from simulations (4000 simulations for every such sub-trajectory). The log-likelihood is $LL(m,\theta) = \sum_j \sum_i \log\left(Prob_{m,\theta,i}\left(X_{i,j}|X_{i-1,j}\right)\right)$, *n=294* sub-trajectories. For each model, we sought the parameter set that maximizes the log-likelihood (see Supplementary Section 1 for more details). Confidence intervals for the best-fit parameters, as well as for estimates for SnC half-life, were calculated by bootstrapping (selecting mice at random with replacements). Modeling experimental noise (multiplicative noise with amplitude up to 30%) did not affect the best-fit parameters (Supplementary Section 1).

To find parameters for the model that describe both the longitudinal trajectories as well as mouse mortality statistics, we scanned the subset of parameters that fit the mortality distribution of mice. Mortality statistics of WT (C57BL/6J) mice were obtained from the Mouse Phenome Database (*1*). Because mortality in young mice appears to be unassociated with the accumulation of senescent cells (*2*), we only considered deaths that occurred after age one year (which make up 97% of the total deaths in the dataset). We performed a comprehensive scan of values of $\beta_0$ and $\kappa_0$ and constrained $\eta, \epsilon$ to values that give a mean and standard deviation of the simulated mortality distributions that is within 2% of the empirical values for WT mice. The critical SnC level $X_C$ was set at $X_C = 17$, which is the maximal SnC level in the Burd et al dataset. We then calculated maximum likelihood parameters and confidence intervals as described above.

#### Population-level measures.

The mean and variance at time-point *i* are the mean and variance of $\{X_{i,j}\}_{j=1}^{N}$ where *N* is the number of mice. Autocorrelation is the Pearson correlation of the two vectors $\{X_{i,j}\}_{j=1}^{N}, \{X_{i+1,j}\}_{j=1}^{N}$. The measures were calculated in the same manner from model simulations. Typical SnC removal rate (half-life$^{-1}$) for the model at a given age *i* was estimated by $\frac{\beta}{\kappa+\bar{X}_i}\log(2)^{-1}$ where $\bar{X}_i$ is the mean SnC level at age *i* (see

Supplementary Section 4 for discussion of alternative ways to estimate SnC half-life). For the USR model, typical SnC removal rate at age $i$ is $(\beta_0 - \beta_1 i)\log(2)^{-1}$.

### Quantification of senescent cells in mouse lung epithelium

We subjected 3-month-old (young) and 22-month-old (aged) C57BL6 mice to intra-tracheal installation of 1.5 U/kg Bleomycin (Sigma) solution in PBS (or PBS as a control treatment). We euthanized the mice at 14, 18, 24 and 28 days for the young mice and day 14 and day 38 for the aged mice. The quantification of senescent epithelial cells was performed as previously described (*3*) with modification. Lung tissue was chopped into 2-5 mm pieces in HBSS (14025050, Gibco) on ice and incubated in the 5ml dissociation buffer (1mg/ml Collagenase Type IV (C9263, Sigma), 0.1 mg/ml DNase I (10104159001, Roche) in HBSS) at 37°C for 50 minutes. Cells were washed with HBBS and then fixed with 4% PFA for 5 minutes. Post fixation, cells were washed and incubated with X-Gal staining solution for 16 h at 37°C. The X-Gal staining solution consisted of 5mM K3Fe(CN)6, 5mM K4Fe(CN)6x3H2O and 2.5 mM X-Gal (Inalco) in PBS at pH5.5 containing 1mM MgCl2. Post X-Gal staining the cells were fixed with fixation buffer for 30 min at 4°C and washed with permeabilization buffer (00-5223-56, eBioscience, San Diego, CA). The cells were then incubated with PE-conjugated pan-cytokeratin (ab52460, Abcam) and HMGB1 (ab18256, Abcam) antibodies for an hour at 4°C. For visualization of HMGB1 antibody, we used Qdot605 labeled Goat Anti-Rabbit antibody (Q11402MP, ThermoFisher). Before visualization, the cells were stained with DAPI and filtered through a 100 µm membrane. The resulting cells were analyzed by imaging flow-cytometry using ImageStreamX mark II (Amnis, Part of EMD milipore - Merck, Seattle, WA, USA, see Supplementary Information 3 for gating strategy summary). PE staining was collected at channel 3, the DAPI at channel 7 and the Qdot605 at channel 10, in addition to the bright-field images collected at channels 1 and 9. Analysis of the image data was performed using IDEAS 6.2 software. Cells were first gated according to their area (in $\mu m^2$) and aspect ratio (ratio between width and length) of the bright field images, to eliminate debris and aggregates. Then, we gated on focused cells using the gradient RMS (which measures the sharpness quality of an image by using the average gradient of a pixel normalized for variations in intensity levels) and contrast (measures the sharpness quality of an image by detecting large changes of pixel values). Cropped cells were excluded by using the centroid X feature (the number of pixels in the horizontal axis from the upper, left corner of the image to the center of the image mask). To verify that only single cells were analyzed, cells were further gated for single nuclei using the area and aspect ratio of the nuclear image of the DAPI staining. SA-beta-Gal staining was quantified using the Mean pixel(the mean of the background-subtracted pixels) contained in the bright-field image, as previously described(*3*). Staining of pCK was quantified using the Intensity (the sum of the background subtracted pixel values within the image) and the Max pixel (the largest value of the background-subtracted pixels contained in the image) features of the corresponding channels. To quantify nuclear staining of HMGB1 specifically, its intensity only within the nuclear mask (made using Morphology mask on the DAPI staining) was calculated. We first gated for pCK positive then for nuclear HMGB1 negative, SA-β-Gal positive cells to quantitate the senescent cells in lung epithelium. Following the method establishment pCK positive, SA-β-Gal positive cells were considered senescent in further experiments. In total, the mice analyzed were 3-month-olds treated with 1.5U/kg (n=16), 22-month-olds treated with 1.5U/kg (n=13), and 22-month-olds treated with PBS (n=6).

#### Analysis of the bleomycin treatment time series.

We estimated the turnover of senescent cells by calculating the removal time after a perturbation with bleomycin. The removal rate is the slope of the log-linear regression model, which we fit for each experiment (with confidence intervals calculated by bootstrapping). We obtained the response time predicted by the model by bootstrapping and simulating the model after perturbation (see Supplementary Section 4 for details).

#### Estimation of hazard and survival functions.

We fit hazard and survival functions from mortality data by interpolation using the Mathematica (V11.0) function SmoothKernelDistribution and then applying the Mathematica functions HazardFunction and SurvivalFunction. For the mice survival data, the SmoothKernelDistribution was computed with a bandwidth of 80 days.

#### Simulation of *Drosophila* and *C. elegans* survival curves

We simulated the mortality trajectories of *Drosophila* and *C. elegans* using the SR model, by assuming a rapid turnover and saturation $\beta = 1\ hr^{-1}$, $\kappa = 1[au]$, and also set $\epsilon = 1\ [au]^2 hr^{-1}$ where [au] is the mean level of X in young organisms. These parameters correspond to a turnover of X on the order of hours. For *C. elegans*, to fit the survival curve obtained by Stroustrup et al., we set $\eta = 0.07[au]hr^{-1}day^{-1}$ and assumed that death occurs when $X > X_C$ for $X_C = 20[au]$. Similarly, to fit the Mair et al. data, we set the following parameters for the *Drosophila* simulations: $\eta = 0.03[au]hr^{-1}day^{-1}$ and $X_C = 15[au]$, and assumed a baseline mortality of $\ln hazard = -7 day^{-1}$.

### Supplementary Section 1. Stochastic modeling of longitudinal trajectories of SnCs in mice.

#### Model comparison and best-fit parameters.

In this section, we consider different stochastic models for the dynamics of senescent cell (SnC) abundance, denoted by $X$. $X$ is removed at rate $removal$ and produced at rate $production$, and includes a noise term *noise*:

$$\dot{X} = production - removal + noise$$

We modeled four biological processes: (i) SnC production rate can increase with age due accumulation of mutations, telomere damage and other forms of intracellular damage that can trigger cellular senescence (*4*), (ii) SnCs can catalyze their own production by paracrine effects (*5*), (iii) SnC removal can decrease with age due to age-related decline in immune surveillance function (*6*) and (iv) SnCs can slow their own removal, for example by saturating or downregulating their own immune surveillance mechanisms. All combinations of these options (including or discluding i-iv) lead to 2^4=16 different circuits:

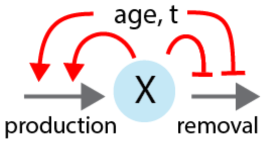

The model that includes all four processes is

$$\dot{X} = (\eta_0 + \eta_1 t)(1 + \eta_2 X) - \frac{\beta_0 - \beta_1 t}{1 + \beta_2 X} X + \sqrt{2\epsilon}\xi_t$$

Where $X$ is SnC abundance, $t$ is time, $\eta_0$ is the initial SnC production rate, $\eta_1$ is the increase in SnC production rate with age, $\eta_2$ is the autocatalysis rate, $\beta_0$ is the initial removal rate, $\beta_1$ is the decrease in removal rate with age, $\beta_2^{-1}$ is the half-way saturation point for removal and $\epsilon$ is the noise amplitude. The most general model has, therefore, 7 parameters.

We defined variants of this model, by all combinations of setting model parameters to zero. A model with no age-related rise in production has $\eta_1$=0, a model with no autocatalysis has $\eta_2$=0, a model with no age related decline in removal has $\beta_1$=0, and a model with no saturation of removal has $\beta_2$=0. The scanned models also include models where there is no removal (i.e. $\beta_0 = \beta_1 = 0$). In this way, we scanned all 1-parameter models, 2-parameter models, etc., up to the full 7-parameter model.

For each model, we scanned parameters to find the parameters that maximize the log-likelihood of the measured longitudinal SnC trajectories (as described in the methods section). We performed the parameter scan for each model in several steps. First a sparse grid was used, in which each parameter was assigned values spaced uniformly on a log scale between $e^{-10}$ to $e^{1}$. Each parameter could also be assigned 0, and in total the number of parameter sets in each grid was at least 50,000 parameter sets. We then used a finer and finer uniform grid within the 95% bootstrapped confidence intervals estimated by the previous sparser scan, until convergence was reached. The scans yielded the following estimates for the log-likelihood of each model (models without time-dependence of parameters (that is, with $\eta_1 = \beta_1 = 0$) show very low likelihood (LL<-550) and are not shown):

| Model # | Model | #Params (best fit) | Saturating removal | Log-likelihood |
|---|---|---|---|---|
| 1 | $\dot{X} = (\eta_0 + \eta_1 t) - \beta_0 X + \sqrt{2\epsilon}\xi_t$ | 4 (4) | no | -535 |
| 2 | $\dot{X} = (\eta_0 + \eta_1 t)(1 + \eta_2 X) - \beta_0 X + \sqrt{2\epsilon}\xi_t$ | 5 (5) | no | -507 |
| 3 | $\boldsymbol{\dot{X} = \eta_0 - (\beta_0 - \beta_1 t)X + \sqrt{2\epsilon}\xi_t (USR\ model)}$ | **4 (3)** | **no** | **-500** |
| 4 | $\dot{X} = (\eta_0 + \eta_1 t) - (\beta_0 - \beta_1 t)X + \sqrt{2\epsilon}\xi_t$ | 5 (3) | no | -500 |
| 5 | $\dot{X} = \eta_0(1 + \eta_2 X) - (\beta_0 - \beta_1 t)X + \sqrt{2\epsilon}\xi_t$ | 5 (3) | no | -500 |
| 6 | $\dot{X} = (\eta_0 + \eta_1 t)(1 + \eta_2 X) - (\beta_0 - \beta_1 t)X + \sqrt{2\epsilon}\xi_t$ | 6 (3) | no | -500 |
| 7 | $\dot{X} = \eta_0 - \frac{\beta_0 - \beta_1 t}{1 + X\beta_2} X + \sqrt{2\epsilon}\xi_t$ | 5 (5) | yes | -483 |
| 8 | $\dot{X} = \eta_0(1 + \eta_2 X) - \frac{\beta_0 - \beta_1 t}{1 + X\beta_2} X + \sqrt{2\epsilon}\xi_t$ | 6 (6) | yes | -479 |
| 9 | $\boldsymbol{\dot{X} = (\eta_0 + \eta_1 t) - \frac{\beta_0}{1 + X\beta_2} X + \sqrt{2\epsilon}\xi_t\ (SR\ model)}$ | **5 (4)** | **yes** | **-475** |
| 10 | $\dot{X} = (\eta_0 + \eta_1 t) - \frac{\beta_0 - \beta_1 t}{1 + X\beta_2} X + \sqrt{2\epsilon}\xi_t$ | 6 (4) | yes | -475 |
| 11 | $\dot{X} = (\eta_0 + \eta_1 t)(1 + \eta_2 X) - \frac{\beta_0}{1 + X\beta_2} X + \sqrt{2\epsilon}\xi_t$ | 6 (6) | yes | -473.7 |
| 12 | $\dot{X} = (\eta_0 + \eta_1 t)(1 + \eta_2 X) - \frac{\beta_0 - \beta_1 t}{1 + X\beta_2} X + \sqrt{2\epsilon}\xi_t$ | 7 (6) | yes | -473.7 |

**Table S1.** Maximum log-likelihood scores for models. Number of parameters in parenthesis the effective number of parameters needed for maximum likelihood, since in some models some of the best-fit parameters are negligible

For the unsaturated removal models ($\beta_2$=0), we also tested an additional model with a quadratic autocatalysis term $\eta_3 X^2$ (because the linear autocatalysis term otherwise could be factored into the removal rate). With this term, the result model has 7 parameters, and has a maximum log-likelihood of -489.

As can be seen from the table above, the best-fit models all have saturating removal ($\beta_2 > 0$). The best-fit model shows removal rates of days in young mice and weeks in old mice. The USR model, which is the best-fit model with unsaturated removal ($\beta_2 = 0$), has a poorer likelihood, and show much longer half-life for SnC. The minimal best-fit model is the 4-parameter SR model with linearly increasing production and constant maximal removal rate, which we rewrite as:

$$\dot{X} = \eta t - \frac{\beta X}{\kappa + X} + \sqrt{2\epsilon}\xi_t$$

(where $\beta = \frac{\beta_0}{\beta_2}, \kappa = \beta_2^{-1}$). Adding temporal changes in the parameters $\kappa, \epsilon$ does not yield improved fits. The best-fit parameters for the SR model are shown in Table S2, with confidence intervals obtained by bootstrapping using the mice individuals in the dataset. SnC levels are given in arbitrary units [au] such that young mice have a mean level of 1[au] in the luciferase dataset.

| Parameter | Mean | SE | 5%CI | 95%CI |
|---|---|---|---|---|
| $\boldsymbol{\eta}$ | $4.2 \cdot 10^{-4}[au]day^{-2}$ | $0.5 \cdot 10^{-4}$ | $3.5 \cdot 10^{-4}$ | $5 \cdot 10^{-4}$ |
| $\boldsymbol{\beta}$ | $0.27\ day^{-1}$ | 0.04 | 0.21 | 0.32 |
| $\boldsymbol{\kappa}$ | $1.1\ [au]$ | 0.3 | 0.7 | 1.8 |
| $\boldsymbol{\epsilon}$ | $0.14[au]^2 day^{-1}$ | 0.02 | 0.1 | 0.18 |

**Table S2.** Best fit parameters for SR model.

These parameters provide a good fit for the mean ($\frac{\chi^2}{N} = 1.27, N = 10$), autocorrelation ($\frac{\chi^2}{N} = 0.63$), and skewness ($\frac{\chi^2}{N} = 0.79$) of the data. The fit for the standard deviation has a higher error ($\frac{\chi^2}{N} = 3.3$), due to the extremely low standard error of the standard deviation at week 16, which is a clear outlier compared with week 8 and week 24. When excluding this point (week 16), we get a good fit for the standard deviation as well ($\frac{\chi^2}{N} = 1, N = 9$), as well an improved fit for the mean ($\frac{\chi^2}{N} = 0.94$) and skewness ($\frac{\chi^2}{N} = 0.37$), and a similar fit for autocorrelation ($\frac{\chi^2}{N} = 0.67$).

The next best-fit minimal model, without saturation, is the 3-parameter model highlighted in Table S1. This model is called unsaturated removal (USR model). The best fit parameters for this model are the following: $\beta_0 = 0.07 \pm 0.018\ day^{-1}$, $\beta_1 = 0.044 \pm 0.01\ day^{-1}year^{-1}$, $\epsilon = 0.22 \pm 0.03\ [au]^2 day^{-1}$.This model, with its best-fit parameters, describes the data poorly (orange lines in Fig 2C, $\frac{\chi^2}{N} = 7.18$ or $\frac{\chi^2}{N} = 6.15$ without week 16 for mean SnC). These parameters are also the best fit parameters for all models without saturation where $\beta_1 > 0$. Here is the full list of best-fit parameters for all models:

| Model # | $\boldsymbol{\eta_0}\ [au]day^{-2}$ | $\boldsymbol{\eta_1}[au]day^{-1}$ | $\boldsymbol{\eta_2}\ [au]^{-1}$ | $\boldsymbol{\beta_0}\ [day]^{-1}$ | $\boldsymbol{\beta_1}\ [day]^{-2}$ | $\boldsymbol{\beta_2}\ [au]^{-1}$ | $\boldsymbol{\epsilon}\ [au]^2 day^{-1}$ |
|---|---|---|---|---|---|---|---|
| **1** | 0.004 | 0.0001 | 0 | 0.019 | 0 | 0 | 0.1 |
| **2** | 0.004 | 0.00006 | 0.6 | 0.03 | 0 | 0 | 0.14 |
| **3** | 0 | 0 | 0 | 0.07 | 0.044 | 0 | 0.22 |
| **4** | 0 | 0 | 0 | 0.07 | 0.044 | 0 | 0.22 |
| **5** | 0 | 0 | 0 | 0.07 | 0.044 | 0 | 0.22 |
| **6** | 0 | 0 | 0 | 0.07 | 0.044 | 0 | 0.22 |
| **7** | 0.15 | 0 | 0 | 0.8 | 0.0009 | 0.52 | 0.19 |

| | | | | | | | |
|---|---|---|---|---|---|---|---|
| **8** | 0.15 | 0 | 0.04 | 0.9 | 0.0009 | 0.4 | 0.11 |
| **9** | 0 | 0.00042 | 0 | 0.25 | 0 | 1.1 | 0.14 |
| **10** | 0 | 0.00042 | 0 | 0.25 | 0 | 1.1 | 0.14 |
| **11** | 0.35 | 0.00055 | 0.04 | 0.9 | 0 | 1.1 | 0.2 |
| **12** | 0.35 | 0.00055 | 0.04 | 0.9 | 0 | 1.1 | 0.2 |

**Table S3.** Best fit parameters for models specified in Table S1.

### Robustness of parameter estimation to experimental noise.

We next tested whether the estimate of the best-fit parameters is robust to experimental noise. We modeled experimental noise as lognormally-distributed multiplicative noise with mean 1 and standard deviation $\sigma$. We repeated the procedure to calculate the log-likelihood of the data (Methods), except that we multiplied the starting point and ending point of each simulated sub-trajectory by a random number sampled from the noise distribution. We then found the best-fit parameters and confidence intervals as described above. The maximal log-likelihood for the SR model, when considering experimental noise with $\sigma = 0.2$ (20% noise), is improved compared to no noise (-465 compared with -475), and we find the following best-fit parameters:

| Parameter | Mean | SE | 5% CI | 95% CI |
|---|---|---|---|---|
| $\boldsymbol{\eta}$ | $4 \cdot 10^{-4}[AU]day^{-2}$ | $0.7 \cdot 10^{-4}$ | $2.4 \cdot 10^{-4}$ | $6 \cdot 10^{-4}$ |
| $\boldsymbol{\beta}$ | $0.28\ day^{-1}$ | 0.055 | 0.17 | 0.47 |
| $\boldsymbol{\kappa}$ | $1.6[AU]$ | 0.4 | 0.6 | 2.7 |
| $\boldsymbol{\epsilon}$ | $0.13[AU]^2 day^{-1}$ | 0.02 | 0.1 | 0.18 |

**Table S4.** Best fit parameters for SR model, assuming 20% noise.

The resulting best-fit parameters are similar to the best-fit parameters without considering experimental noise, and lead to the same conclusions regarding SnC dynamics, turnover rate, saturation and slowdown. Similar results are found with noise amplitudes of $\sigma = 0.05, 0.1, 0.3$. Therefore, the present parameters are robust to experimental noise in the longitudinal SnC measurements.

### Best-fit parameters adjusted for mortality.

We sought to find the best-fit parameters for the SR model $\dot{X} = \eta t - \frac{\beta X}{\kappa + X} + \sqrt{2\epsilon}\xi_t$, that describe both the longitudinal trajectories and mice mortality statistics. For this purpose, we scanned the values of $\beta$ and $\kappa$ and constrained $\eta, \epsilon$ so the mean and standard deviation of the mortality-time distribution will be within 2% of the data for the mortality distribution of WT (C57BL/6J) mice obtained from the Mouse Phenome Database (*1*) (see Methods). We set mortality as described in the Methods section.

In a simple realization of the connection between SnC and mortality, one can assume that death occurs when SnC level exceed$X_C$. We set $X_C = 17[au]$, which is the maximal observed SnC level in the normalized TBL data. The best-fit maximum likelihood parameters are:

| Parameter | Mean | SE | 5% CI | 95% CI |
|---|---|---|---|---|
| $\boldsymbol{\eta}$ | $2.3 \cdot 10^{-4}[au]day^{-2}$ | $0.25 \cdot 10^{-4}$ | $1.9 \cdot 10^{-4}$ | $2.9 \cdot 10^{-4}$ |
| $\boldsymbol{\beta}$ | $0.15\ day^{-1}$ | 0.022 | 0.12 | 0.2 |
| $\boldsymbol{\kappa}$ | $0.5[au]$ | 0.1 | 0.4 | 0.7 |
| $\boldsymbol{\epsilon}$ | $0.16[au]^2day^{-1}$ | 0.02 | 0.14 | 0.2 |

**Table S5.** Best fit parameters for SR model, adjusted for mortality.

The mortality distribution computed with these best-fit parameters agrees with the mouse mortality distribution (p>0.2 under both Kolmogorov-Smirnov and Anderson-Darling tests). The log-likelihood for the best-fit parameters for the longitudinal SnC data is -490. When excluding the outlier point (week 16), these parameters provide a good fit for the mean ($\frac{\chi^2}{N} = 0.84$, N=9), skewness ($\frac{\chi^2}{N} = 0.28$), and autocorrelation ($\frac{\chi^2}{N} = 0.46$), though the fit for the variance is worse than the parameters of Table S2 ($\frac{\chi^2}{N} = 2.9$).

While these best-fit parameters lead to similar conclusions regarding SnC turnover rate, saturation and slowdown, they are different from the best-fit parameters that are not constrained by mortality (Table S2). Specifically, while noise amplitude $\epsilon$ is similar to the original best-fit value, the other parameters are smaller, leading to higher noise compared with feedback strength. The reason for this may be slightly different survival curves for mice grown in different locales, and/or that the requirement that death occurs at a critical SnC level may be unrealistic compared with a more gradual dependence of mortality on SnC. Such gradual dependence allows for lower noise compared with feedback (Supplementary Section 2).

### Mathematical modelling of the bystander effect of SnC production

Several recent studies suggest that SnC can be produced as a result of direct induction by neighboring SnCs (“bystander effect”) as a result of paracrine interactions (*7–10*). The injection of a small number of senescent cells can spread cellular senescence in host tissues (*11*), providing *in vivo* evidence for the bystander effect.

The contribution of the bystander effect to SnC dynamics can be modelled in several ways. The first and most straightforward is that SnC production rate includes a term that is proportional to SnC abundance $\eta_2 X$. This assumes that each SnC generates more SnCs at a rate $\eta_2$, leading to exponential propagation of SnCs in the tissue. As can be seen in Table S1, this mechanism is not essential to explain the data of Burd et al, because models that include such a bystander effect term do not outperform models without the bystander effect. In the most complex model: $\dot{X} = (\eta_0 + \eta_1 t)(1 + \eta_2 X) - \frac{\beta_0}{1+X\beta_2^{-1}} X + \sqrt{2\epsilon}\xi_t$, the contribution of the bystander effect $\eta_2$ is negligible in the best fit parameter set.

Another possibility is that the bystander effect is non-linear. For example, it may be that SnCs only catalyze their own formation when they are at a high concentration. This nonlinearity can create a local tipping point, where SnCs are induced at a rate higher than their removal once they exceed a certain concentration.

Lastly, it may be that the bystander effect is limited to a specific bounded region within a tissue, or that the paracrine induction of SnCs weakens after near neighbors turn senescent. Future work can address this by considering the spatial dynamics of SnCs in tissues.

### Supplementary Section 2. Analytical properties of the SR model

In this section, we derive various analytical properties of the SR model.

The SR model equation is:

$$\dot{X} = \eta t - \frac{\beta X}{X + \kappa} + \sqrt{2\epsilon}\xi_t \qquad [1]$$

One can write this using a potential U(X,t):

$$\dot{X} = -\frac{d}{dx}U(X,t) + \sqrt{2\epsilon}\xi_t \qquad [2]$$

Where the potential is

$$U(X) = (\beta - \eta\mathrm{t})X - \beta\kappa\log(\kappa + X) \qquad [3]$$

#### Quasi-stationary distribution and statistical properties of SR model

Due to the rapid turnover relative to the organism lifetime, we can use a quasi-steady-state approximation (main text Figure 1B). At quasi-steady state:

$$X_{ST} = \frac{\eta\kappa t}{\beta - \eta t} [4]$$

In a quasi-steady state approximation, the stationary distribution is given by the Boltzmann distribution:

$$Prob[X = x] \propto e^{-\frac{U(x)}{\epsilon}} \qquad [5]$$

Therefore:

$$Prob[X = x] \propto e^{-\frac{(\beta - \eta\mathrm{t})x}{\epsilon}}(\kappa + x)^{\frac{\beta\kappa}{\epsilon}} \qquad [6]$$

With the normalization factor, the distribution is:

$$Prob[X = x] = \frac{e^{-\frac{x(-\eta\mathrm{t}+\beta)}{\epsilon}-\frac{(-\eta\mathrm{t}+\beta)\kappa}{\epsilon}}\kappa^{-1-\frac{\beta\kappa}{\epsilon}}\left(-\frac{(\eta-\beta)\kappa}{\epsilon}\right)^{\frac{\epsilon+\beta\kappa}{\epsilon}}(x+\kappa)^{\frac{\beta\kappa}{\epsilon}}}{\mathrm{Gamma}[1+\frac{\beta\kappa}{\epsilon}, -\frac{(\eta\mathrm{t}-\beta)\kappa}{\epsilon}]} [7]$$

We can use this distribution to approximate the mean SnC abundance:

$$\langle X \rangle = \int_{x=0}^{\infty} x\, Prob[X = x]dx = \frac{\kappa\eta\mathrm{t} + \epsilon + \frac{e^{\frac{\kappa(\eta\mathrm{t}-\beta)}{\epsilon}}\epsilon}{\mathrm{ExpIntegralE}[-\frac{\kappa\beta}{\epsilon}, \frac{\kappa(-\eta\mathrm{t}+\beta)}{\epsilon}]}}{\beta - \eta\mathrm{t}} [8]$$

For parameters in the relevant range: $\frac{e^{\frac{\kappa(\eta\mathrm{t}-\beta)}{\epsilon}}\epsilon}{\mathrm{ExpIntegralE}[-\frac{\kappa\beta}{\epsilon}, \frac{\kappa(-\eta\mathrm{t}+\beta)}{\epsilon}]} \ll \kappa\eta t + \epsilon$ this yields:

$$\langle X \rangle \approx \frac{\kappa\eta t + \epsilon}{\beta - \eta\mathrm{t}} [9]$$

For example, for parameters similar to the best fit mouse parameters, $\beta = 0.2, k = 1, \epsilon = 0.5$, the analytical solution of Eq 8 and its approximation Eq 9 are nearly identical, as shown in Figure S1.

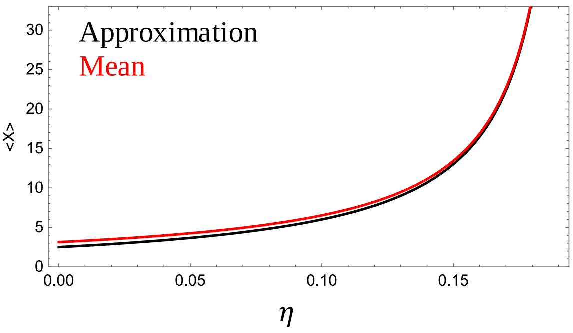

**Figure S1. Analytical approximation of Eq 9 is similar to Eq 8 over the relevant range for η.** Equations 8,9 were simulated given the physiologically relevant parameters $\beta = 0.2, k = 1, \epsilon = 0.5$, and varying η (corresponding to the dynamics of aging under the SR model). Both equations predict similar mean SnC level <X>.

The variance is estimated as:

$$\langle X^2 \rangle = \frac{\epsilon\left(\kappa\beta + \epsilon - \frac{e^{\frac{\kappa(\eta t-\beta)}{\epsilon}}\left(e^{\frac{\kappa(\eta t-\beta)}{\epsilon}}\epsilon + \kappa p \mathrm{ExpIntegralE}\left[-\frac{\kappa\beta}{\epsilon}, \frac{\kappa(-\eta t+\beta)}{\epsilon}\right]\right)}{\mathrm{ExpIntegralE}\left[-\frac{\kappa\beta}{\epsilon}, \frac{k(-\eta t+\beta)}{\epsilon}\right]^2}\right)}{(\eta-\beta)^2} \approx \frac{\kappa\beta + \epsilon^2}{(\eta t-\beta)^2} \qquad [10]$$

The distribution of SnC in the SR model is skewed to the right (Eq. 7) quantitatively matching the skewness observed in the mouse data (main text Figure 2EF).

### Derivation of Gompertz mortality in the SR model

We model mortality as the first time when $X > X_C$ (we later discuss other cases). Thus, death time is a first-passage time of the SR model variable X. To estimate the hazard rate (probability of death per unit time), we apply the Kramer approximation for the first passage time (*12*, *13*):

$$h \approx \frac{\sqrt{U''(X_{ST})U''(X_C)}}{2\pi} e^{-\frac{U(X_C)-U(X_{ST})}{\epsilon}}$$

Where the effective potential U is given by Eq.3. For the Gompertz law(*14–18*) to hold, one needs $\frac{U(X_C)-U(X_{ST})}{\epsilon}$ to decrease linearly with time, so that $h \approx e^{\alpha t}$.

The exponent of the hazard rate in the SR model indeed shows the required linearity in time:

$$-\frac{U(X_C)-U(X_{ST})}{\epsilon} = \frac{(\kappa + X_C)\eta t - X_C\beta + \kappa\beta \cdot \mathrm{Log}\left[\frac{(\kappa+X_C)(\beta-\eta t)}{\kappa\beta}\right]}{\epsilon} \qquad [8]$$

The curvature around steady-state is $U''(X_{ST}) = \frac{(\beta-\eta)^2}{\kappa\beta}$. We also denote $U''(X_C) = \omega_{max}$ as the (unknown) curvature around the critical threshold. We thus find that:

$$h \approx \frac{\sqrt{\omega_{max}}}{2\pi}(\kappa + X_C)^{\frac{\kappa\beta}{\epsilon}}(\kappa\beta)^{-\frac{\kappa\beta}{\epsilon}-0.5}(\beta-\eta t)^{\frac{\kappa\beta}{\epsilon}+1} e^{\frac{(\kappa+X_C)\eta t - X_C\beta}{\epsilon}} \qquad [9]$$

The hazard rises exponentially with time as $e^{\alpha t}$ with $\alpha = \frac{(\kappa+X_C)\eta}{\epsilon}$. The model also shows a deceleration in the rise of the hazard rate at very old ages (when $\eta t \approx \beta$), due to the prefactor $(\beta-\eta t)^{\frac{\kappa\beta}{\epsilon}+1}$, as observed for empirical hazard (*19*, *20*). Note that this approximation begins to be inaccurate when $\eta t > \beta$, and simulations of the full SR model are needed to compute the hazard curve at old ages.

Models without SnC turnover (models with $\beta_0 = \beta_1 = 0$) do not yield a mortality distribution that follows the Gompertz law, but, instead, an inverse Gaussian distribution (*19*, *21*).

### Near-exponential rise of mean SnC in the SR model

The dependence of mean SnC on time in the SR model is similar to an exponential, in the sense that it matches the second order Padé approximation of an exponential:

$$k \cdot e^{\frac{\eta_1}{\beta}t} \approx k\frac{1+\frac{\eta_1}{\beta}t}{1-\frac{\eta_1}{\beta}t} = \frac{k\beta + k\eta_1 t}{\beta - \eta_1 t} \approx \frac{\kappa\eta_1 t + \epsilon}{\beta - \eta_1 t} \approx \langle X \rangle \quad [10]$$

The last equality holds when $k\beta \approx \epsilon$. This is the case for the best-fit mouse parameters. More generally, the resemblance to the exponent holds when $k, \beta, \epsilon$ are of similar magnitude (as can be shown by simulation). A semi-logarithmic plot of mean SnC as a function of age in the SR model appears nearly linear (model 9, Table S1), whereas the USR with best fit parameters (model 3, Table S1) deviates from linearity.

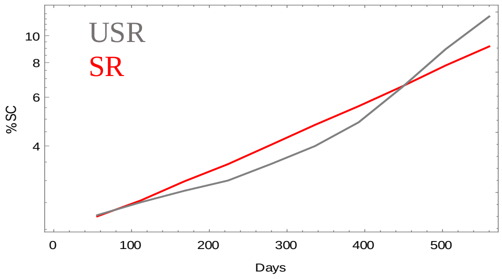

**Figure S1. Mean SnC increases approximately exponentially for the SR model and super-exponentially for the USR model.** The USR model (model 3 in table S1) the SR model (model 9 in table S1) were simulated with their best-fit parameters (table S4). The increase in mean SnC is approximately exponential for the SR model and super-exponential for the USR model. This indicates that the SR model provides a nearly exponential increase in SnC levels as observed.

#### Gompertz law holds to a good approximation under a general dependence of mortality on SnC level.

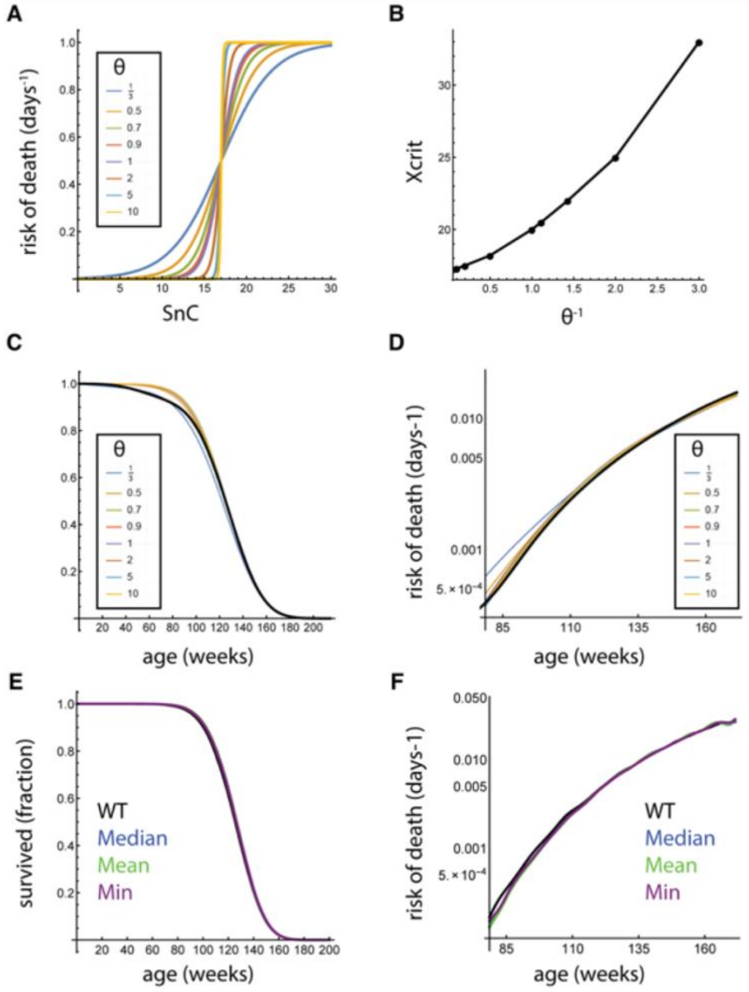

**Figure S3. Gompertz law holds under a general dependence of mortality on SnC level.** (A) We tested whether the Gompertz law holds in the SR model under the more general assumption that probability of death increases with SnC level $X$ as a logistic function with steepness $\alpha$ and half-way point $X_C$. In the case of $\alpha \gg 1$ this converges to a step function in which death occurs when crossing a threshold $X_C$. (B) Using the best-fit parameters provided in Section 1, Table S5, we show that for each steepness $\alpha$ we can choose an appropriate $X_C$ such that both the hazard curves (C) and the mortality curves (D) fit the observed mouse mortality statistics well (black curves in panels C-F). (EF) We tested whether the Gompertz law holds under the more general assumption that death occurs only when the average/median/minimal number of SnC over a longer time period (30 days) exceeds a thresholds $X_C$. We show that with appropriate choices of $X_C$ ($X_C = 17$ for mean, median, $X_C = 15$ for min), both the hazard curves (E) and the mortality curves (F) fit the observed mouse mortality statistics well.

In the previous section we showed that the SR model leads to the Gompertz law, assuming that death occurs when SnC level $X$ crosses a critical level $X_C$. Here we show by simulation that this conclusion also holds under a more general dependence of death on SnC level. We model probability of death as a function of X as a sigmoidal increasing function with different degrees of steepness. The probability of death is described using a family of logistic functions, in which steepness depends on a parameter $\alpha$:

$$Prob(death, x) = \frac{1}{1 + e^{-\alpha(X - X_C)}}$$

The larger $\alpha$, the steeper the dependence of probability of death on $X$, around the half-way point $X_c$ (Figure S3A, similar results are found using Hill functions). For each of the above choices of $\alpha$ one can choose an appropriate $X_C$ so that mortality distributions are very similar to the distribution observed for mice (Figure S3B-D), keeping the other model parameters constant at their values in Table S5. This indicates that an exponentially rising increase of mortality with age (Gompertz law) plus slowdown at old ages is a property that holds even when assuming a graded probability of death as a function of SnC level.

We also tested whether the Gompertz law holds if we assume that death occurs when SnC exceed a critical threshold for a period of time (30 days). We tested the cases where death occurs when average SnC, median SnC or minimal SnC over 30 days exceed a critical threshold. For each of the above we can choose an appropriate $X_C$ so that mortality distributions are very similar to the distribution observed for mice (Figure S3EF), suggesting the generality of Gompertz law for this condition as well.

### Supplementary Section 3. Gating strategy employed to identify senescent cells.

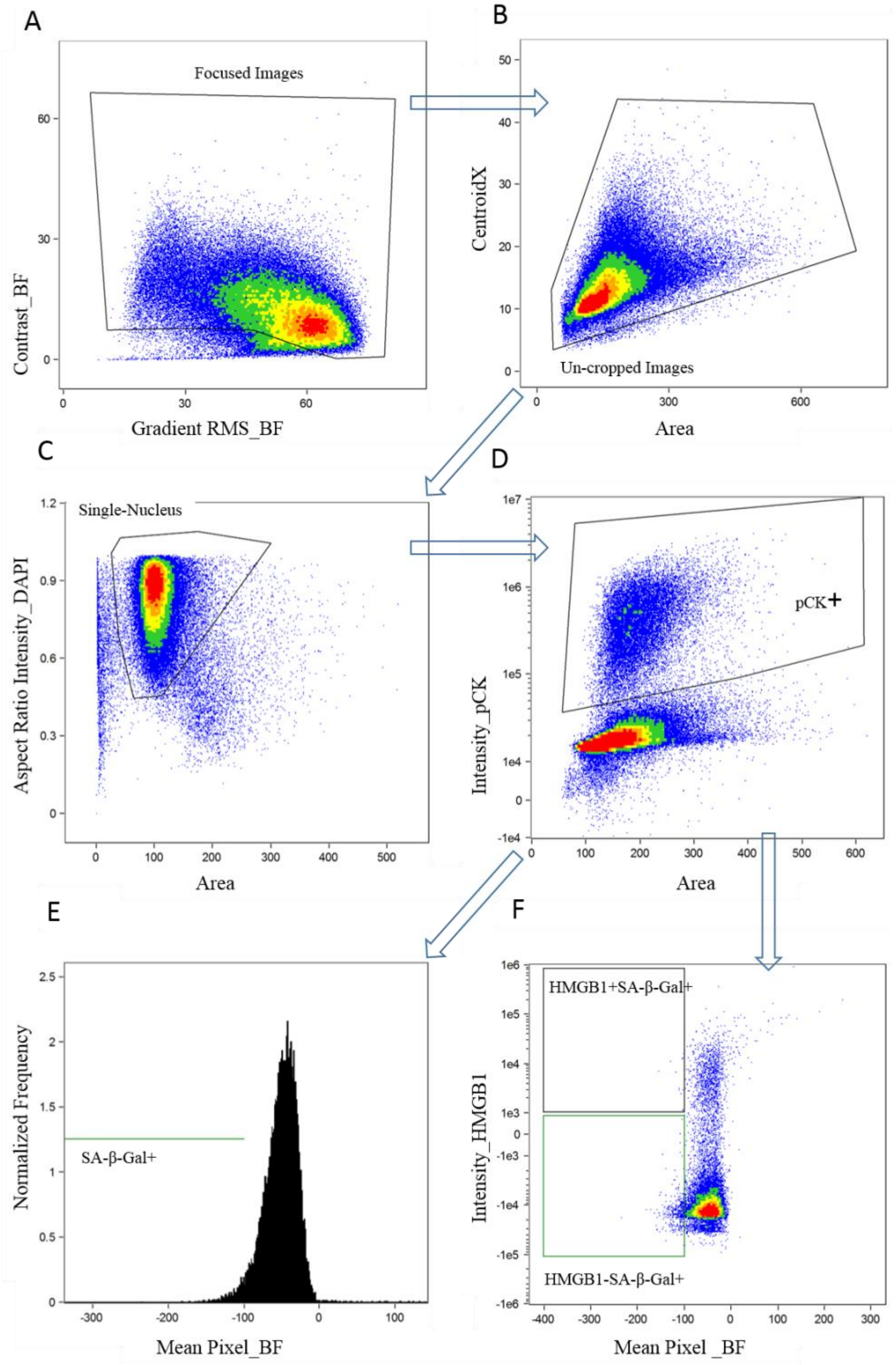

**Figure S4. Gating strategy employed to identify SA-β-Gal+ or SA-β-GAL+HMGB1- cells in the lung epithelium using ImageStreamX.** Gating strategy employed to identify SA-β-Gal+ or SA-β-GAL+HMGB1- cells in the lung epithelium using ImageStreamX. **(A)** Focused images were gated using Gradient Root Mean Square and contrast (both measure the sharpness quality of an image by detecting large changes of pixel values in the image) of the Bright Field (BF) channel. **(B)** Uncropped images were gated out of Focused images using CentroidX (distance of a cell from the left side of the acquired image) and Area in square microns of the bright field image, **(C)** Single nucleus cells were selected using area and aspect ratio (normalized for intensity) of the DAPI staining out of uncropped images, **(D)** pan cytokeratin (pCK) positive cells were selected using intensity and the bright field area out of single nucleus cells, **(E)** Out of pCK positive cells, SA-β-Gal+ cells were selected using the mean pixel (the mean of the background-subtracted pixels contained in the input mask) of the Bright Field, where SA-β-Gal+ cells are darker and show lower values as previously described (*22*). **(F)** SA-B-Gal+ and HMGB1- cells were gated using HMGB1 intensity and mean pixel of the Bright Field channel.

### Supplementary Section 4. Estimation of SnC removal rate.

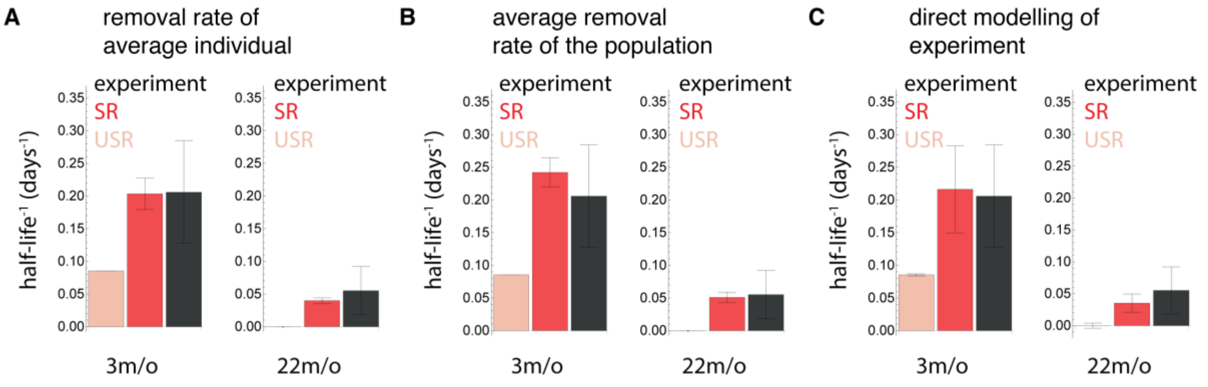

**Figure S5. Estimation of SnC removal rates by different methods yields similar results.** SnC removal rate of the SR production-removal model that was fit to the longitudinal trajectories was estimated by different methods (red bars): (A) Removal rate of the average individual: $\frac{\beta}{\kappa+\bar{X}_\iota}\log(2)^{-1}$, (B) Average removal rate of the population: $\overline{\frac{\beta}{\kappa+X_\iota}}\log(2)^{-1}$, and (C) Modeling of bleomycin perturbation experiments. Model predictions were compared with estimation from bleomycin perturbation experiments (Figure 2 in the main text). The model shows similar predictions across estimation methods (red, mean and SE were calculated by bootstrapping according to the number of mice in each bleomycin experiment). Removal rate for the next best-fit model that does not have saturation (USR mode, gray) was estimated in a similar manner.

In this section, we compare three ways to use the SR model to simulate the bleomycin experiment and compute the half-life of SnC levels. We find that the three ways provide very similar results.

The SR production-removal model for SnC dynamics:

$$\dot{X} = \eta t - \frac{\beta X}{\kappa + X} + \sqrt{2\epsilon}\xi_t$$

has a per-SnC removal rate of $\frac{\beta}{\kappa+X}$, where $X$ is the total SnC abundance. Therefore, a perturbation in which SnC abundance is changed by $\delta$ ($\delta \ll X$) will decay with a half-life$^{-1}$ of $\frac{\beta}{\kappa+X+\delta}\log(2)^{-1} \approx \frac{\beta}{\kappa+X}\log(2)^{-1}$.

Since SnC abundance $X$ is heterogeneous in the population, the model predicts that SnC removal rate will also be heterogeneous. We considered three ways to estimate the typical removal rate at a given age $i$.

1. The removal rate of the average individual: $\frac{\beta}{\kappa+\bar{X}_\iota}\log^{-1} 2$ where $\bar{X}_\iota$ is the mean SnC level at age $i$. This calculation predicts a SnC half-life of about $5 \pm 1$ days in young (3-month-old) mice and a SnC half-life of about $25 \pm 6$ days in old (22-month-old) mice.
2. The average removal rate of the population: $\overline{\frac{\beta}{\kappa+X_\iota}}\log^{-1} 2$.This calculation predicts SnC half-life of about $4 \pm 1$ days in young (3-month-old) mice and a half-life of about $20 \pm 5$ days in old (22-month-old) mice.

Both estimates yield similar predictions for SnC dynamics that are consistent with the half-life estimated from the perturbation experiments (Figure 2 in the main text, Figure S5AB).

3. We also modeled the experiments directly, by bootstrapping SnC levels $X_j$ at age $i$ and simulating the ODE: $\dot{x} = \eta i - \frac{\beta x}{\kappa + X_j}$.We set an initial post-bleomycin level $x\ (5\ days) = X_j + 50[au]$, representing a perturbation that increases lung epithelium SnC level to $50[au]$ after 5 days (the conclusions are not dependent on the exact initial value). For each time point of the bleomycin experiment $T_j \in \{T_1 \dots T_n\}$ we sampled $X_j$ and simulated the above ODE, and measured the level of $x(T_j)$. We then estimated SnC removal rate in the same manner as for the bleomycin time-series (Figure S5C, Figure 2C). This method yields an estimated removal rate that is similar to that predicted by methods 1, 2.

We estimated removal time in the same manner for the USR model: $\dot{X} = \eta - (\beta_1 - \beta_0 t)X + \sqrt{2\epsilon}\xi_t$. Since the best-fit USR model predicted a slightly negative removal rate for 22-month-old mice, we set its removal rate at that age to zero. All estimation methods show that the USR model provides a poorer prediction for the SnC half-life (Figure S5).

### Supplementary Section 5. SR model for SnC dynamics.

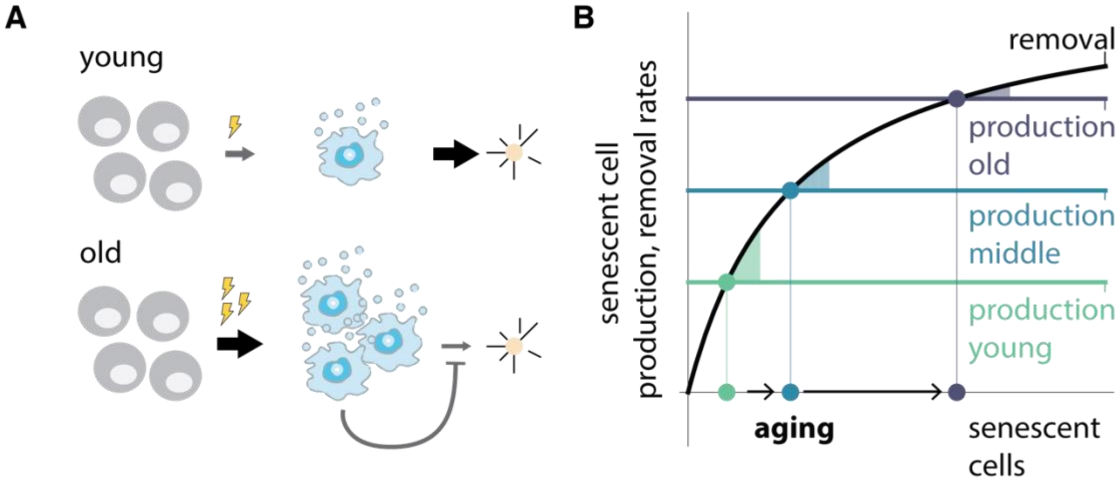

**Figure S6. Saturating removal and increasing production lead to accelerated SnC abundance with age and persistent SnC fluctuations.** (A) With age, production rate increases, SnC levels rise and increasingly saturate their own removal mechanism, further amplifying their rate of increase. (B) Removal and production rates plotted as a function of SnC abundance. Steady-state occurs when production and removal curves cross (dots). As production rises, SnC levels accelerate to higher abundance. At old ages, the spare capacity for removal (the distance between production and removal curves, shaded regions) shrinks, so that SnC perturbations last longer, leading to persistent fluctuations. These fluctuations cause non-genetic variations between individuals that last for long times.

### Supplementary Section 6. Predicted effect of senolytic drug regimes

We employed the SR model to simulate the effects of drugs that eliminate SnCs, known as senolytic drugs (*9*, *23–29*). Due to toxicity concerns, it may be desirable to establish regimes of low dose and large temporal spacing for these drugs (*25*). We computed the effect of regiments of senolytic drug treatments as a function of their intake frequency and efficacy.

To model the effect of a senolytic drug, we consider $X_{SENSITIVE}$ to be the number of senescent cells that are sensitive to the drug and $X_{INSENSITIVE}$to be the number of senescent cells that are not sensitive to the drug ($X_{INSENSITIVE}$ may also represent other forms of damage that impacts aging besides senescent cells). Overall, the level of $X$ is: $X = X_{SENSITIVE} + X_{INSENSITIVE}$.We assume that a fraction $\zeta$of SnC production and noise goes to make $X_{SENSITIVE}$, and the rest $(1-\zeta)$ goes to $X_{INSENSITIVE}$:

$$\dot{X}_{SENSITIVE} = \zeta\eta t - \frac{\beta X_{SENSITIVE}}{X+\kappa} + \sqrt{2\zeta\epsilon}\xi'_t$$

$$\dot{X}_{INSENSITIVE} = (1-\zeta)\eta t - \frac{\beta X_{INSENSITIVE}}{X+\kappa} + \sqrt{2(1-\zeta)\epsilon}\xi''_t$$

All parameters are the same as for mouse (Table S5), and death occurs when $X > X_C = 17[AU]$. To model the effect of a drug given every $\phi$ days, we define a term $f(\phi,\alpha,t)$ which represents the killing rate of the drug:

$$f(\phi,\alpha,t) = \begin{cases} \alpha & if\ t\ mod\ \phi = 0 \\ 0 & otherwise \end{cases}$$

Thus $\phi$ is the period at which the drug (time between doses) is taken and $\alpha$ is the drug efficacy (rate of killing sensitive cells). Thus:

$$\dot{X}_{SENSITIVE} = \zeta\eta t - f(\phi,\alpha,t)X_{SENSITIVE} - \frac{\beta X}{X_{SENSITIVE}+\kappa} + \sqrt{2\zeta\epsilon}\xi_t$$

We set $\zeta$ in mice by assuming that the pharmacogenetic experiments of Baker et al.(*30*), that used a genetic construct to destroy p16-expressing cells, mimic the effect of a drug with a maximal SnC removal rate of $\alpha_{MAX} = 1day^{-1}$. In these experiments, $\phi = 1day$ and mean lifespan of the mice increased by 25%. This allows calibrating $\zeta$, providing $\zeta \approx 0.25$. Simulating the model for different $\phi,\alpha$ gives[38] the following dependence of predicted life extension on the time between doses and efficacy per dose:

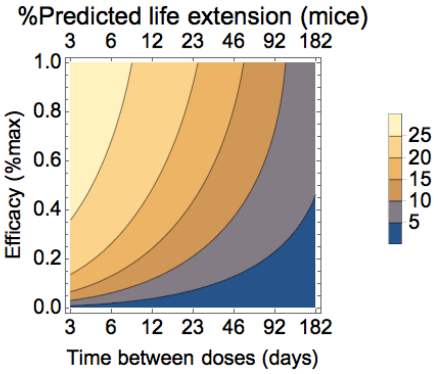

The parameters used for the simulations are the mortality-adjusted best-fit parameters (Table S5). We can thus see that effective treatment can be as infrequent as the SnC turnover time (~month in old mice) even for a drug that kills only a fraction of the sensitive SnC.

### Supplementary Section 7. Human aging dynamics and the SR model.

To study human mortality curves we used the Swedish life table for 2009, obtained from http://www.lifetable.de. This dataset is standard for demographic research on mortality statistics (*16, 31–33*). We model the hazard using the hazard from the SR model plus an age-independent extrinsic mortality, $h = h_{SR} + h_{extrinsic}$, with $h_{extrinsic} = 0.4 \cdot 10^{-3} year^{-1}$ estimated from mean mortality at age 20 (Makeham term (*34*)). As discussed in the main text, taking the mortality-adjusted mouse parameters in the SR model (Table S5) and adjusting $\eta$ by ~1/60 is sufficient to obtain a good fit to the human mortality data, without adjusting additional parameters (Figure 3).

To test the plausibility of these parameters, we compared the model predictions to measurements of SnC concentrations in various human tissues using different methods (Table S6).

| Tissue (Human) | Method | N (number of subjects) | SnC accumulation slope [1/year] |
|---|---|---|---|
| **T-cells (*35*)** | p16 mRNA content | 156 | 3.7±0.4% |
| **Bone marrow stromal cells (hMSC) (*36*)** | SA-beta-gal | 16 | 3.5±1% |
| **Renal arteries (*37*)** | %p16-positive nuclei | 30 | 2±1% |
| **Renal glomureli (*37*)** | %p16-positive nuclei | 36 | 2.1±1.1% |
| **Renal interstitium (*37*)** | %p16-positive nuclei | 35 | 4±1.5% |
| **Renal tubules (*37*)** | %p16-positive nuclei | 39 | 4±0.9% |
| **Corneal epithelium (*38*)** | IHC staining of p16 | 12 | 3.5±1.2% |
| **Cartilage chondrocytes (*39*)** | SA-beta-gal | 15 | 2.0%±0.5% |

**Table S6.** SnC accumulation slope for various human tissues.

Data was taken from the figures in the indicated studies. Data points with very small SnC abundance (log value less than -1) were capped at -1. Since we only seek to model SnC levels at adulthood, we used data points after age 20 (results are unaffected by using data at all ages). Slope was estimated using regression on log data, with bootstrapped error bars. The measurements show an approximately exponential increase of SnC levels with a rate of 2%-4% per year. This rate of increase is consistent with the present values for $\kappa$ and $\beta$ for mice, which provide an increase of ~2.2% per year in total body SnC. We conclude that $\kappa$ and $\beta$ mouse parameters are a plausible choice for human SnC accumulation rates.

### Supplementary Section 8. SR model and mortality interventions in *Drosophila*

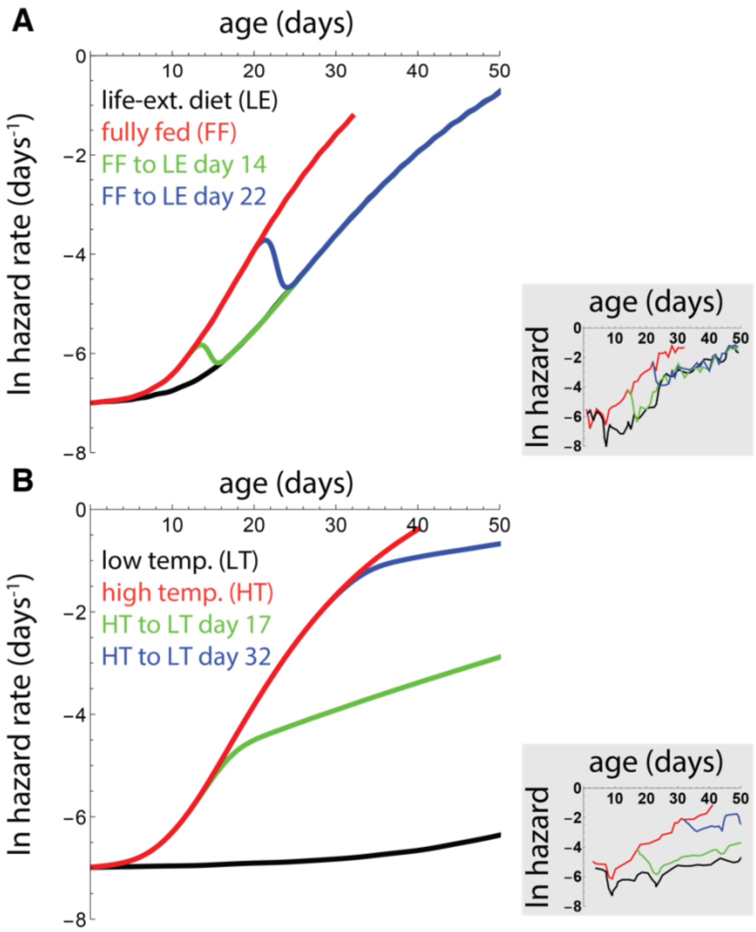

**Figure S7. Rapid turnover and saturating removal can explain the effect of mid-life interventions on *Drosophila* mortality.** (A) The SR model with saturation and rapid turnover of damage ($\beta = 1\ hr^{-1}$, $\kappa = 1[AU]$, $\epsilon = 1\ [AU]^2 hr^{-1}$,$X_C = 15[AU]$), and a slow increase in damage production ($\eta = 0.03[AU]hr^{-1}day^{-1}$) can explain the hazard curve of fully fed *Drosophila* (red curve). A slower rate of increase in damage production $\eta = 0.02[AU]hr^{-1}day^{-1}$ can explain the hazard curve of *Drosophila* under dietary change (LD, black curve). An instantaneous change in $\eta$ leads to rapid reversal of mortality rates (green and blue lines), explaining the observation of Mair et al. (insets). (B) The SR model with a slow rate of increase in damage production $\eta = 0.005[AU]hr^{-1}day^{-1}$ can explain the hazard curve of *Drosophila* raised at 18C (low temperature, black line). A change in the underlying rate of increase in damage production leads to a change in the slope of the hazard, as observed by Mair et al (insets).

In a classic paper, Mair et al. (*40*) measured the effect of two lifespan-extending interventions in *Drosophila*, lifespan-extending diets (LE) and temperature change, when applied at mid-adulthood (*40*, *41*). They found that the interventions had different effects on lifespan: (i) LE led to rapid switches in mortality rate, and (ii) changing temperature affected the slope of the mortality rate.

These results can be explained by the SR model with rapid turnover (Figure S7). A relatively rapid turnover for *Drosophila* means turnover on the order of minutes to hours. We therefore set $\beta = 1\ hr^{-1}$, $\kappa = 1[AU]$, and $\epsilon = 1\ [AU]^2 hr^{-1}$. To fit the survival curve for fully fed flies obtained by Mair et al., we set $\eta = 0.03[AU]hr^{-1}day^{-1}$, and death when $X > X_C$ with $X_C = 15[AU]$. Flies undergoing LE are fit by a lower value, $\eta = 0.02[AU]hr^{-1}day^{-1}$. Note that the purpose here is to demonstrate that the SR model can capture the behavior of the data, and not to provide accurate estimates for the parameters (the data is insufficient to pin down the parameters).

The LE dietary intervention can be explained by assuming that it changes any of the model parameters. For example, LE can change $\eta$ in a reversible manner (Figure S7A), and hence affect the rate of damage production $p$. In this case, changing diet leads to damage production $p(t) = \eta_1 t,$ where $\eta_1$ is the rate of increase in damage production of the current diet. The rapid turnover of damage rapidly reverts the mortality rates when diet changes (Figure S7A). More generally, LE may change any parameter of the SR model, including removal rate β, as long as the effect on the parameter is reversible.

On the other hand, the temperature intervention can be explained by assuming that it affects an underlying damage accumulation rate that sets $\eta$ (Figure S7B), that is, temperature multiplies $\frac{dp}{dt}$. Changing temperature at age $t$' therefore leads to damage production $p(t) = \eta_0 t' + \eta_1(t - t')$, where $\eta_0$ was the previous rate of increase in damage production and $\eta_1$ the rate after temperature change. This intervention affects the slope of increase in mortality rate with age, but does not revert the mortality rates (Figure S7B).

### Supplementary Section 9. SR model and scaling of *C. elegans* survival curves

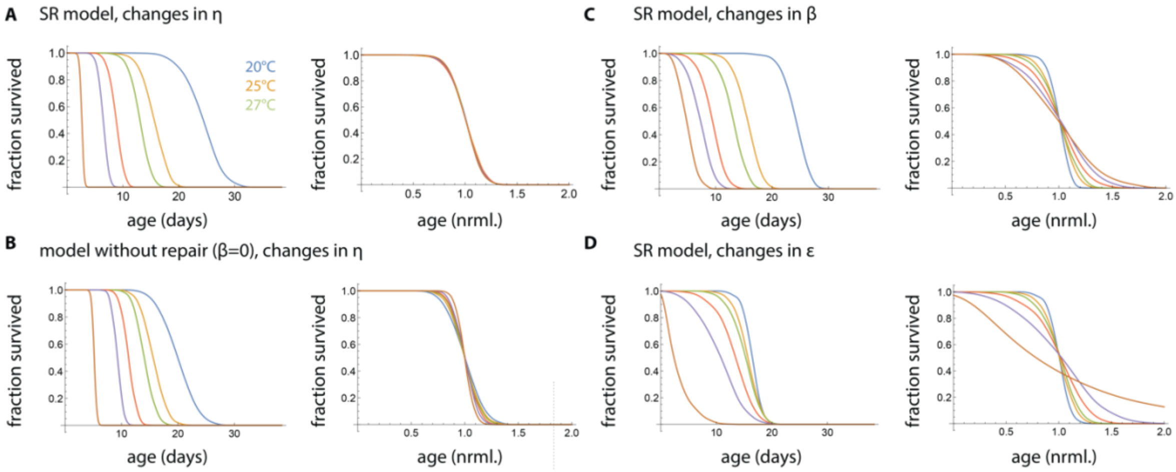

**Figure S8. SR model with rapid turnover can explain the temporal scaling of survival curves in *C. elegans*.** (A) The SR model with saturation and rapid turnover of damage ($\beta = 1\ hr^{-1}$, $\kappa = 1[AU]$, $\epsilon = 1\ [AU]^2 hr^{-1}$,$X_C = 20[AU]$), and a slow increase in damage production ($\eta = 0.07[AU]hr^{-1}day^{-1}$) can explain the survival distribution of *C. elegans* raised at room temperature (25C). Changes in damage production rate $\eta$, by multiplying it by a factor $\lambda$ ($\lambda \in \{0.6, 1, 1.25, 2, 3, 10\}$), lead to lifespan distributions that are temporally scaled, despite a variation of over an order of magnitude in mean lifespan (right panel is age normalized by mean lifespan). (B) A model without damage turnover that was fit to the same survival curve ($\beta = 0$, $\eta = 0.006[AU]hr^{-1}day^{-1}$,$\epsilon = 0.037\ [AU]^2 hr^{-1}$) does not show time scaling of survival curves. Time scaling is also not observed for the SR model for changes in $\beta$ (C) or $\epsilon$ (D). Each line represents the simulation of 1000 individuals, time scaling was confirmed according to the statistical criteria of Stroustrup et al. (*42*).

In an elegant study, Stroustrup et al. measured lifespan distributions under various conditions that affect mean lifespan, such as temperature, nutrition and mutations (*42*). The aging distributions were different in each condition, but for most perturbations, the curves collapsed to the same curve when normalizing time by a scaling constant. Stroustrup et al. also found that transient temperature interventions shift the lifespan distribution, and concluded that risk of death is determined by a single stochastic process, with a single effective rate constant which is altered by the interventions.

We tested whether the SR model is consistent with these findings, and if so, which parameter is likely to be affected by the lifespan-changing conditions. Note that in this case the form of the damage (the meaning of X in the model) is unknown.

We first estimated a set of physiologically reasonable parameters for the SR model for *C. elegans*. Since *C. elegans* grown at room temperature lives for ~2 weeks on average, a relatively rapid turnover means turnover on the order of minutes to hours. We therefore set $\beta = 1\ hr^{-1}$, $\kappa = 1[AU]$, and $\epsilon = 1\ [AU]^2 hr^{-1}$. To fit the survival curve obtained by Stroustrup et al., we set $\eta = 0.07[AU]hr^{-1}day^{-1}$ and death when $X > X_C$ with $X_C = 20[AU]$.

We find that changes in the parameter $\eta$ of the SR model can explain the scaling observations of Stroustrup et al. We find that changes in $\eta$ (i.e. multiplying $\eta$ by λ, λ∈{0.6,1,1.25,2,3,10}) result in lifespan distributions that are different, but collapse nearly perfectly onto a single distribution when time is normalized by the mean lifespan (Figure S8A).

Importantly, time scaling requires rapid turnover. For example, a model without rapid turnover ($\beta = 0$) does not show time scaling of survival curves (Figure S8B). Similarly, scaling breaks down when turnover time is comparable to lifespan.

We find that changes in parameters other than $\eta$ do not provide time scaling of survival curves (Figure S8C,D). We therefore conclude that the SR model can explain time scaling of survival curves, and identify $\eta$ as the parameter that is modulated by the interventions.

Generally, the scaling of $\eta$ needed to fit survival curves in different interventions is inversely proportional to the mean lifespans. For example, for the mutation interventions specified by Stroustrup et al., we find a scaling of $\lambda = 0.75$ for *daf-2(e1368)*, $\lambda = 2$ for *daf-16(mu86)* and $\lambda = 1.2$ for *hsf-1(sy441)*. The effects of other time-scaling interventions on $\eta$ can be similarly calibrated from Supplementary Table 2 in Stroustrup et al. (*43*)

Stroustrup et al. also reported interventions that do not show time scaling. These perturbations may be explained by changes that affect additional parameters besides $\eta$. For example, survival of the *eat-2 (ad1116)* mutant is well fit by multiplying $\eta$ by $\lambda = \frac{2}{3}$ and multiplying noise $\epsilon$ by $\frac{3}{2}$. Survival of the*nuo-6 (qm200)* mutant is well fit by multiplying $\eta$ by $\lambda = \frac{1}{2}$ and multiplying noise $\epsilon$ by 2. These findings suggest that it would be intriguing to further explore experimentally how these interventions affect stochastic damage dynamics.
